# A simple model-based approach to inferring and visualizing cancer mutation signatures

**DOI:** 10.1101/019901

**Authors:** Yuichi Shiraishi, Georg Tremmel, Satoru Miyano, Matthew Stephens

## Abstract

Recent advances in sequencing technologies have enabled the production of massive amounts of data on somatic mutations from cancer genomes. These data have led to the detection of characteristic patterns of somatic mutations or “mutation signatures” at an unprecedented resolution, with the potential for new insights into the causes and mechanisms of tumorigenesis.

Here we present new methods for modelling, identifying and visualizing such mutation signatures. Our methods greatly simplify mutation signature models compared with existing approaches, reducing the number of parameters by orders of magnitude even while increasing the contextual factors (e.g. the number of flanking bases) that are accounted for. This improves both sensitivity and robustness of inferred signatures. We also provide a new intuitive way to visualize the signatures, analogous to the use of sequence logos to visualize transcription factor binding sites.

We illustrate our new method on somatic mutation data from urothelial carcinoma of the upper urinary tract, and a larger dataset from 30 diverse cancer types. The results illustrate several important features of our methods, including the ability of our new visualization tool to clearly highlight the key features of each signature, the improved robustness of signature inferences from small sample sizes, and more detailed inference of signature characteristics such as strand biases and sequence context effects at the base two positions 5’ to the mutated site.

The overall framework of our work is based on probabilistic models that are closely connected with “mixed-membership models” which are widely used in population genetic admixture analysis, and in machine learning for document clustering. We argue that recognizing these relationships should help improve understanding of mutation signature extraction problems, and suggests ways to further improve the statistical methods.

Our methods are implemented in an R package **pmsignature** (https://github.com/friend1ws/pmsignature) and a web application available at https://friend1ws.shinyapps.io/pmsignature_shiny/.

**Author Summary:** Somatic (non-inherited) mutations are acquired throughout our lives in cells throughout our body. These mutations can be caused, for example, by DNA replication errors or exposure to environmental mutagens such as tobacco smoke. Some of these mutations can lead to cancer.

Different cancers, and even different instances of the same cancer, can show different distinctive patterns of somatic mutations. These distinctive patterns have become known as “mutation signatures”. For example, C *>* A mutations are frequent in lung caners whereas C *>* T and CC *>* TT mutations are frequent in skin cancers. Each mutation signature may be associated with a specific kind of carcinogen, such as tobacco smoke or ultraviolet light. Identifying mutation signatures therefore has the potential to identify new carcinogens, and yield new insights into the mechanisms and causes of cancer,

In this paper, we introduce new statistical tools for tackling this important problem. These tools provide more robust and interpretable mutation signatures compared to previous approaches, as we demonstrate by applying them to large-scale cancer genomic data.

## Introduction

Cancer is a genomic disease. As we lead a life, DNA within our cells acquires random somatic mutations, mainly caused by DNA replication errors and exposures to mutagens such as chemical substances, radioactivities and inflammatory reactions. Although most mutations are harmless (called “passenger mutations”), a small portions of mutations at some specific sites in cancer genes (“driver mutations”) affect cell growth, causing autonomous proliferation, tissue invasion, and contributing to oncogenesis [1]. Cancer genome studies typically focus on identifying driver mutations, to help understand the mechanism of cancer development. However, passenger mutations can also yield important information, because they often show patterns (“mutation signatures”) which can provide insights into the forces that cause somatic mutations. For example, classical studies of mutation patterns revealed that C *>* A mutations are abundant in lung cancers in patients with smoking history, and these are caused by benzo(a)pyrene included in tobacco smokes [2]. Also, C *>* T and CC *>* TT mutations are abundant in ultraviolet-light-associated skin cancers, and these are caused by pyrimidine dimers as a result of ultraviolet radiation [3].

The potential for classical studies to yield insights into somatic mutation processes were limited in several ways. Due to limited sequencing throughputs, most classical studies focused on a few cancer genes, such as TP53, where high mutation frequencies could be expected. They then contrasted mutation pattern profiles among different cancer types, aggregating mutations across multiple individuals within the same cancer type to yield sufficient mutations for analysis. However, since many of the mutations in cancer genes are driver mutations causing cell proliferation, the resultant mutation profiles are a biased representation of the underlying mutation process. Furthermore, the paucity of mutation data made it effectively impossible to assess variation in mutation patterns among individuals.

Recent advances in high-throughput sequencing provide new opportunities to investigate sample-by-sample mutation signatures in an unbiased way using genome-wide somatic mutation data. For example, a large-scale study using 21 breast cancer samples identified an association of C *>* [AGT] mutations at TpC sites, which was later proved to be caused by APOBEC protein family [4–6], and a novel phenomenon called *kataegis* [7]. Moreover, a landmark study of 7,034 primary cancer samples, representing 30 different cancer classes, has provided the first large-scale overview of mutation signatures across a large number of cancer types [8]. This has lead to great hopes that detection of novel mutation signatures and associated mutagens can lead to identification of novel mutagens and prevention of cancer.

To make the most of these opportunities requires the development of efficient and effective statistical methods for analyzing mutation signatures in vast amounts of somatic mutation data. Current statistical approaches [9, 10], are an excellent starting point, and have helped generate the new insights noted above. However, we argue here that these existing methods have two important limitations, caused by the fact that they use an unconstrained model for each “mutation signature.” First, although using an unconstrained model might appear to be a good thing in terms of flexibility, in practice it can actually reduce flexibility, because the price of using an unconstrained model is that one must limit the domain of mutation signatures considered. For example, most recent analyses of mutation signatures consider only the immediate flanking 5’ and 3’ bases of each substitution to be part of the signature, even though it is known that more distal bases – and particularly the next flanking base on each side – can contain important contextual information [11]. These recent analyses take this approach because, in the unconstrained model, incorporating the more distal bases into the signature very substantially increases the number of parameters, making estimated mutation signatures unstable. Secondly, and just as important, the unconstrained model means that each signature is a probability distribution in a high-dimensional parameter space, which can make signatures difficult to interpret.

In this paper, we present a novel probabilistic approach to mutation signature modelling that addresses these limitations. In brief, we first simplify the modelling of mutation signatures by decomposing them into separate “mutation features”. For example, the substitution type is one feature; flanking bases are each another feature. We then exploit this decomposition by using a probabilistic model for signatures that assumes independence across features. This approach substantially reduces the number of parameters associated with each signature, greatly facilitating the incorporation of additional relevant sequence context. For example, our approach can incorporate both the two bases 3’ and 5’ of the substitution, and transcription strand biases using only 18 parameters per signature, compared with 3071 parameters per signature with current approaches. We demonstrate the benefits of this simplification in data analyses. These benefits include: more stable estimation of signatures from smaller samples, refinement of the detail and resolution of many mutation signatures, and, possibly, identification of novel signatures.

Assuming independence among features in a signature may initially seem unnatural. However, its use here is analogous “position weight matrix models” which have been highly successful for modelling transcription factor binding motifs. Indeed, an important second contribution of our paper is to provide intuitive visual representations for mutation signatures, analogous to the “sequence logos” used for visualizing binding motifs. Finally, we also highlight the close connection between mutation signature models and the “mixed-membership models”, also known as”admixture models” [12] or “latent Dirichlet allocation” models [13] that are widely used in population genetics and document clustering applications. These connections should be helpful for future elaboration of computational and statistical methods for cancer mutation signature detection.

Software implementing the proposed methods is available in an R package **pmsignature** (**p**robabilistic **m**utation signature), at https://github.com/friend1ws/pmsignature. The core part of the estimation process is implemented in C++ by way of the Rcpp package [14], which enables handling millions of somatic mutations from thousands of cancer genomes using a standard desktop computer. In addition, a web-based application of our method is available at https://friend1ws.shinyapps.io/pmsignature_shiny/.

## 1 Result

### New model for mutation signatures

The term “mutation signature” is used to describe a characteristic mutational pattern observed in cancer genomes. Such patterns are often related to carcinogens (e.g., frequent C > A mutations in lung cancers with smoking histories).

What constitutes a mutational pattern varies among papers. The simplest approach is to consider 6 possible mutation patterns, corresponding to 6 possible substitution patterns (C*>*A, C*>*G, C*>*T, T*>*A, T*>*C, T*>*G; the original base is often fixed to C or T to remove redundancy of taking complementary strands). However, in practice we know that DNA context of a substitution is often important, and so it is common to go the next level of complexity, and include the immediate 5’ and 3’ flanking bases in the mutation pattern. This results in 96 (6 × 4 × 4) patterns. Further incorporating the strand (plus or minus) of each substitution extends this to 192 patterns [8, 9].

Mathematically, mutation signatures have previously been characterized using an unconstrained distribution over mutation patterns [9, 10]. Thus, if the number of mutation patterns considered is *M* then each mutation signature is characterized by a probability vector of length *M* (which must sum to 1, so *M −* 1 parameters). A problem with this approach is that it requires a large number of parameters per mutation signature. As noted above, even accounting only for immediately flanking bases gives *M* = 96. Furthermore, *M* increases exponentially if we try to account for additional context: to take account of up to *-n* bases 5’ and 3’ to the mutated site (henceforth referred to as the *n* position and the +*n* position, respectively) gives *M* = 6 ×4^2*n*^. Having a large number of parameters per signature causes two important problems: i) estimates of signature parameters can become statistically unstable; ii) signatures can become difficult to interpret.

The first contribution of this paper is to suggest a more parsimonious approach to modelling mutation signatures, with the benefit of producing both more stable estimates and more easily-interpretable signatures. In brief, we substantially reduce the number of parameters per signature by breaking each mutation pattern into “features”, and assuming independence across mutation features. For example, consider the case where a mutation pattern is defined by the substitution and its two flanking bases. We break this into three features (substitution, 3’ base, 5’ base), and characterize each mutation signature by a probability distribution for each feature (which, by our independence assumption, are multiplied together to define a distribution on mutation patterns). Since the number of possible values for each feature is 6, 4, and 4 respectively this requires 5 + 3 + 3 = 11 parameters instead of 96 *−* 1 = 95 parameters. Furthermore, extending this model to account for *±n* neighboring bases requires only 5 + 6*n* parameters instead of 6×4^2*n*^ *–* 1. For example, considering *±*2 positions requires 17 parameters instead of 1535. Finally, incorporating transcription strand as an additional feature adds just one parameter, instead of doubling the number of parameters.

Since the aim of a mutation signature is, in some ways, to capture dependencies among features, the independence assumption may seem counter-intuitive. However, the idea is exactly analogous to the use of a “position weight matrix” (PWM) to represent motifs in sequence data. In this analogy, a motif is analogous to a mutation signature, and each location in the motif is analogous to a “feature”. Just as we use a probability vector for each feature, a PWM defines a probability vector for each location in a motif, and these probabilities at each location can be multiplied together to yield a probability distribution on sequences. Even though a PWM cannot capture complex higher-order dependencies, some of which likely do exist in practice, it has been a highly successful tool for motif analysis - likely because it can capture the most important characteristics of transcription factor binding sites (that some locations will show strong preference for a particular base, whereas others will not), and also because it can be represented in an easily-interpretable way via sequencing logos [15]. For similar reasons – in addition to the empirical demonstrations we present later – we believe our mutation signature representation will prove useful for mutation signature analysis.

Fig. 1 illustrates the way that our new representation of signatures can simply capture a previously-identified signature [9, 16] and provides an easily-interpretable visualization of the signature that is analogous to sequencing logos [15]. We particularly note how the key elements of this mutation signature are more immediately visually apparent than with visualizations of the full vector of probabilities used by existing approaches.

**Figure 1.**
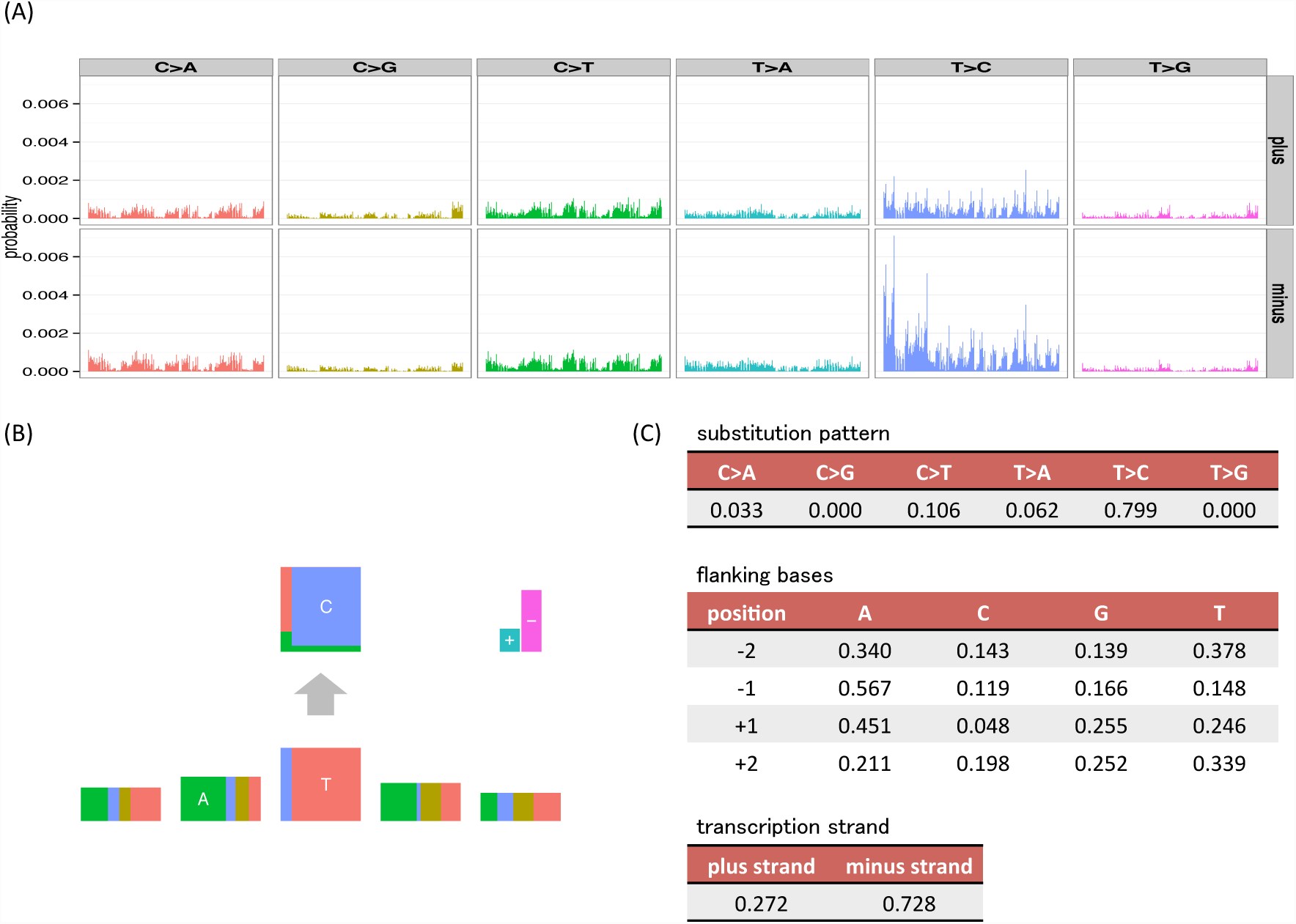
Examples of visualizations and parameter values for the mutation signatures of the unconstrained (full) model and our independent model, where substitution patterns, two 5’ and 3’ bases and transcription strand direction are considered as mutation features. (A) The barplots are divided by 6 substitution patterns and transcription strand direction. In each division, 256 bars show joint probabilities of up to two base 5’ and 3’ bases (ApApNpApA, ApApNpApC, ApApNpApG, ApApNpApT, …, TpTpNpTpT). (B, C) An example mutation signature representation and parameter values from our independent model, where mutation features (substitution patterns, two 5’ and 3’ bases and strand direction) are assumed to be independent (*L* = 6, ***M*** = (6, 4, 4, 4, 4, 2)). In the bottom five rectangles, the width of each box represents the frequencies of bases (A, C, G and T) at the substitution and flanking site. To highlight the most informative flanking sites, the heights of flanking site boxes are scaled by (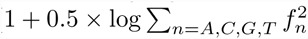, where *f*_*n*_ is the parameter for each base), which can be interpreted as (1 0.5 Rényi entropy) [17]. This is analogous to the information content scaling used in sequencing logos. In the top rectangle, the height of each box represents the conditional frequencies of mutated bases for each original base (C and T). In the upper right, the height of the + box represents the frequencies of mutations in the coding strand (or the plus strand, the sense strand and the untranscribed strand) whose nucleotide sequences directly corresponds to mRNA, whereas the height of – box represents those in the template strand (or the minus strand, the antisense strand, the transcribed strand and the noncoding strand) whose sequences are copied during the synthesis of mRNA.

### An overview of mathematical specification of mutation signatures and the generative model of somatic mutations

Suppose each somatic mutation has *L* mutation features, ***m*** = (*m*_1_, *m*_2_, *…, m*_*L*_), where each *m*_*l*_ can take *M*_*l*_ discrete values. Also, let ***M*** := (*M*_1_, *…, M*_*L*_). For example, when taking account of 6 substitution patterns and *±*2 flanking sites, ***M*** = (6, 4, 4, 4, 4). See S1 Table for other examples.

Suppose we have observed mutations in *I* sampled cancer genomes, and let *J*_*i*_ denote the number of observed mutations in the *i*-th cancer genome. Further, let ***x***_*i,j*_ = (*x*_*i,j,*1_, *…, x*_*i,j,L*_), (*i* = 1, *…, I, j* = 1, *…, J*_*i*_) denote the observed mutation feature vector for the *j*-th mutation of *i*-th cancer genome, where *x*_*i,j,l*_ ∈ {*1, …, M*_*l*_}

Our model assumes that each mutation arose from one of *K* possible mutation signatures. Each cancer sample has its own characteristic proportion of mutations of each signature type (which might depend on lifestyle, genetic differences, etc.). We let *q*_*i,k*_ denote the proportion of signature *k* in sample *i*, so ***q***_*i*_ = (*q*_*i,*1_, *q*_*i,*2_, *…, q*_*i,K*_) ∈ Δ^*K*^, (*i* = 1, *…, I*) where Δ^*S*^ = {(*t*_1_, *…, t*_*S*_) : *t*_*s*_ *≥* 0 *∀s*, 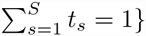} denotes the S-dimensional simplex of non-negative vectors summing to 1. Further, each mutation signature is characterized by parameter vectors ***F***_*k*_ := (***f***_*k,*1_, *…,* ***f***_*k,L*_), where ***f***_*k,l*_ is a probability vector for the *l*-th feature in the *k*-th signature. That is, ***f***_*k,l*_ = (*f*_*k,l,*1_, *…, f*_*k,l,M*_) ∈ Δ^*M*_*l*_^

Our generative model for the observed mutations {***x***_*i,j*_} in each cancer sample can now be described as a two-step process.

1. Generate *z*_*i,j*_ *∼* Multinomial(***q***_*m*_), where *z*_*i,j*_ ∈{1, *…K*} denotes the (unobserved) underlying mutation signature that caused the *j*-th mutation in the *i*-th sample.
2. For each *l*(= 1, *…, L*), generate *x*_*i,j,l*_ *∼* Multinomial(***f***_*z*_*i,j*_,l_). Thus,

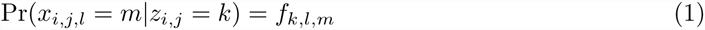

This generative model is summarized in Fig. 2. This model is essentially a “mixed-membership model”, also known as an “admixture model” [12] or “latent Dirichlet allocation” [13]. For example, the membership proportions for each sample are analogous to admixture proportions in an admixture model; the mutation signatures are analogous to populations, and the mutation signature-specific parameters are analogous to population-specific allele frequencies.

**Figure 2.**
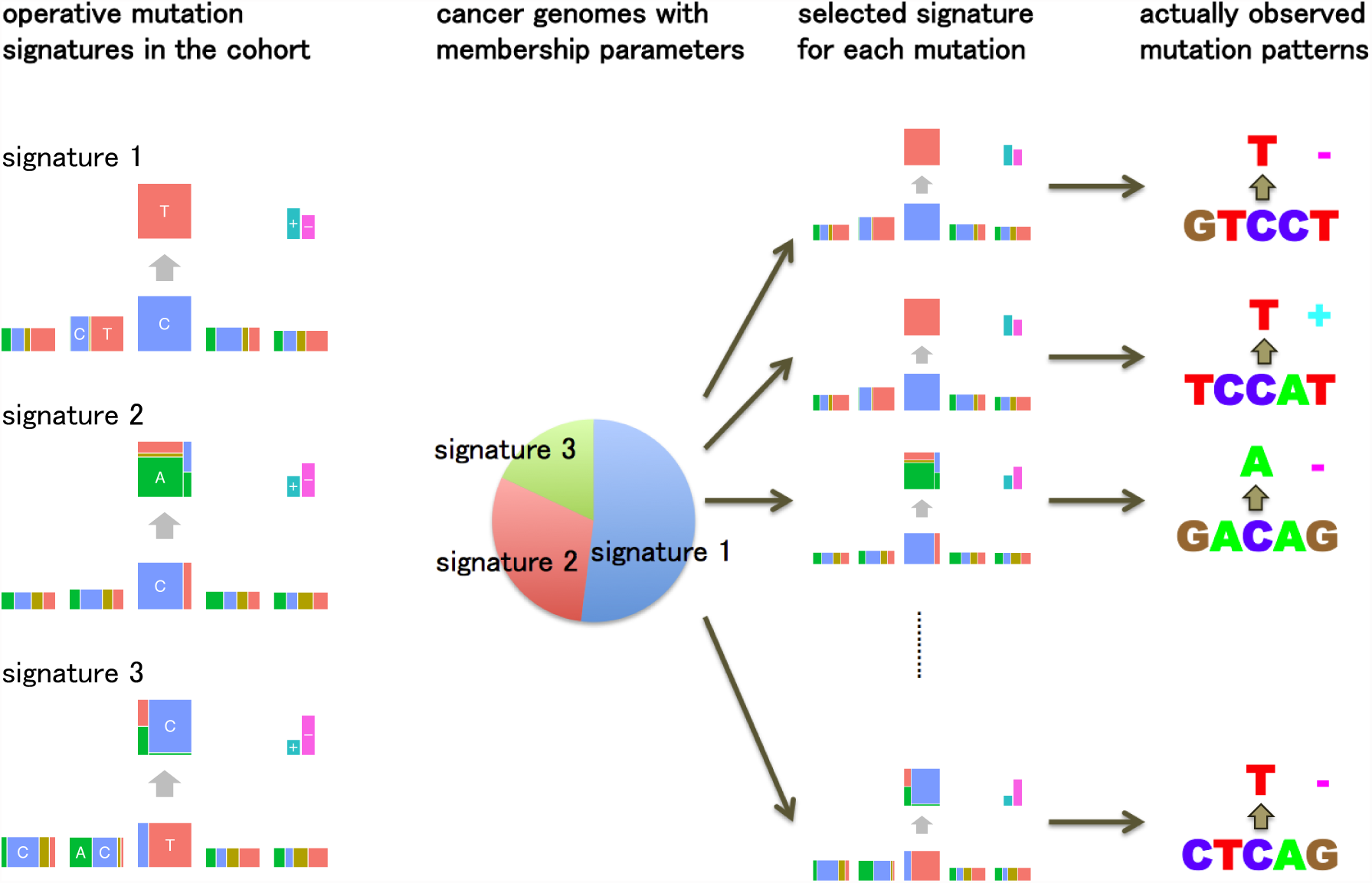
An overview of the generative model of somatic mutations proposed in this paper. Suppose there are three types of mutation sources (mutation signatures) such as ultraviolet, tobacco smoking chemicals and transcription coupled repairs. Each cancer genome has ratios showing which types of mutation sources are contributing to its mutations (membership parameters). The generative model of the pattern of each mutation is: first, one of the mutation signatures is chosen according to the membership parameter. Second, each mutation feature such as substitution patterns and flanking bases is generated by the corresponding multinomial distributions for the selected mutation signature.

The key parameters in this model are the membership proportions for each sample, ***q***_*i*_, and the mutation signature parameters, ***F***_*k*_. We estimate these parameters by maximizing likelihood using an EM algorithm. A simulation study demonstrates that the estimation method can reproduce the mutation signature very accurately provided enough mutations and samples are available (see S1 Text). See Methods for more detailed models, parameter estimation, further discussion on relationships with mixed membership models, how to select *K*, etc.

The intrinsic composition of genome sequence, if unaccounted for, can undesirably influence estimated mutation signatures. For example, since the di-nucleotide CpG is underrepresented in most genomic regions (other than promoters), a signature with substitutions from a C base can have weaker signals of G base at the +1 position. In previous work [10], this background problem was dealt by explicitly incorporating “mutation opportunity” coefficients into the model. Here, to reduce the influences of intrinsic sequence composition on our signature estimates, we introduce a special “background signature”

**{*F***_0_} Δ^*M*_*1*_ *×…×M*_*L*_^, which is designed to capture biases in intrinsic genome sequence composition and is calculated from the composition of consecutive nucleotides of the human genome sequence (See Methods for the detail).

### Robustness experiments using cancer genomes from urothelial carcinoma of the upper urinary tract

Here we compare our new “independent model” for mutation signatures, which assumes independence among mutation features, with the “full model”, which corresponds to existing approaches. We compare mutation signatures obtained by the two approaches and investigate the robustness of each approach by down-sampling experiments.

The data consist of 14717 somatic substitutions collected from a study of 26 urothelial carcinomas of the upper urinary tract (UCUT) [18]. The original study identified a novel mutation signature in these data: T *>* A substitutions at CpTpG sites with a strong transcription strand specificity caused by aristolochic acids (AA).

We consider a mutation pattern to consist of the substitution pattern, the *±*2 flanking bases, and the transcription strand direction. Thus each signature is characterized by 18 parameters in our independent model, and by 3071 parameters in the full model. After analyzing the data with various number of mutation signatures *K*, we selected *K* = 3 signatures for these analyses (see S2 Text).

The inferred APOBEC signature under the independent model shows a clear depletion of G base at the –2 position, which is consistent to the previous study [9] and results in the next subsection (Figs. 3A and 3B). In contrast, for the full model, this tendency is rather mild (Figs. 3C, 3D and S1). The inferred AA mutation signature has no clear characteristics at the –2 position. These results suggest that our independent model has the potential to identify signatures in more detail and with less data than existing approaches based on the full model.

**Figure 3.**
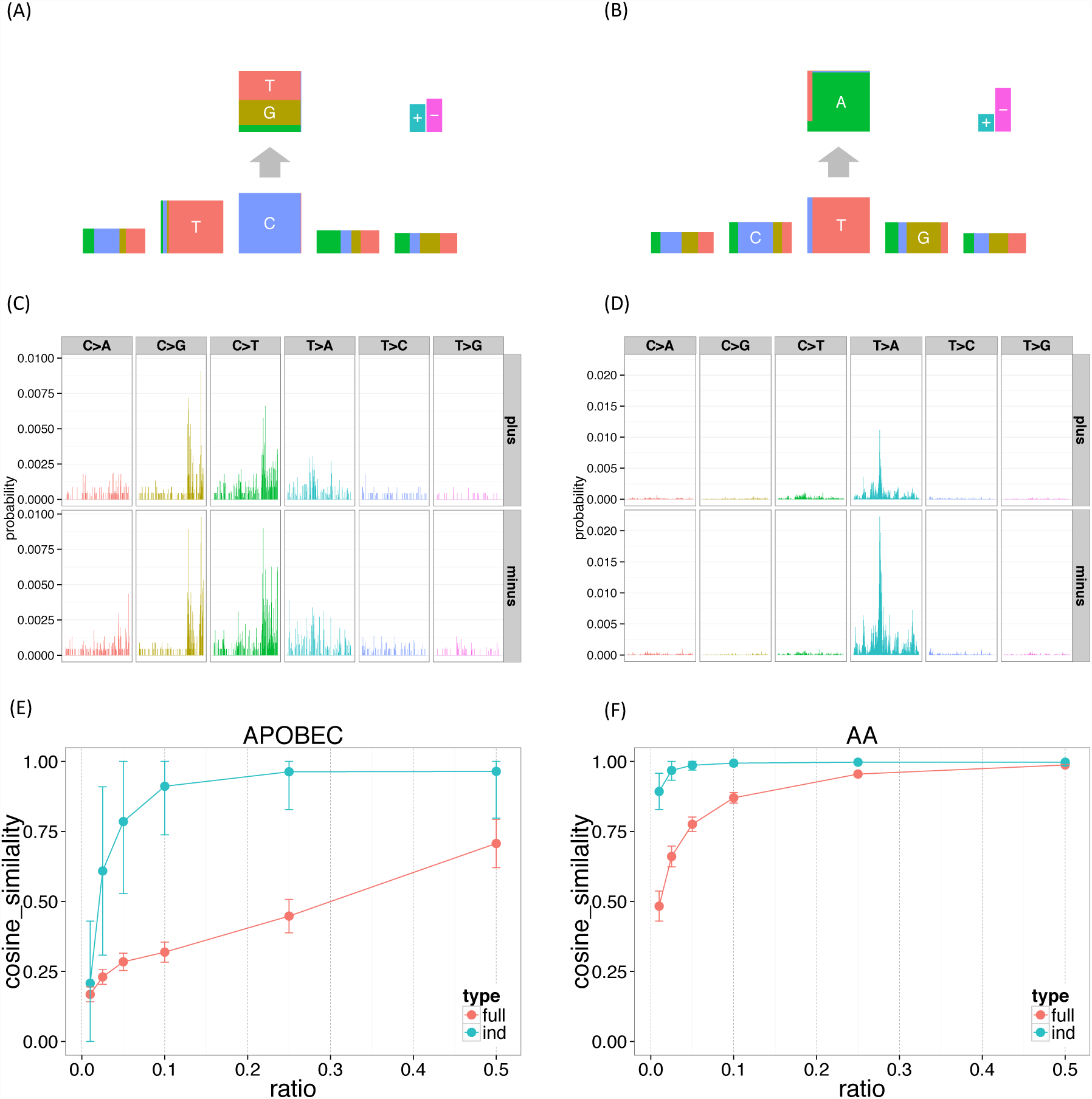
The mutation signatures for the UCUT data, and the results of down-sampling experiments. 3072 elements in the full model mutation signatures were shown divided by 6 substitution patterns and strand directions. (A, B) APOBEC and AA signature for the independent model. (C, D) APOBEC and AA signature for the full model. (E, F) APOBEC and AA signature stability (the mean cosine similarity for each donw-sampling ratio).

To investigate this further we performed down-sampling experiments. Using the mutation signatures obtained using all 14717 substitutions as a gold standard, we assessed performance of the proposed method on down-sampled data consisting of *r*% of the original data, where *r* = (1%, 2.5%, 5%, 10%, 25%, 50%). To measure robustness we used the cosine similarity on the full dimensional vector space, which allows comparison between the full model and the independent model. We repeated each down-sampling experiment 100 times for each model.

The results (Figs. 3E and 3F) confirm that the results of the independent model are substantially more robust to reductions in data size than the full model. Indeed, mutation signatures inferred using the independent model with only 10% of the data remain highly similar to the signatures inferred from the full data; by comparison the full model shows a much larger drop-off in similarity, especially in the APOBEC signature where even using 50% of the data gives a substantial drop-off in similarity. Both methods found the AA signature easier to recover than the APOBEC signature. We believe that this is because the number of T *>* A substitutions at GpTpC sites are far more frequent in this dataset.

### Application to somatic mutation data of 30 cancer types

To provide a more comprehensive practical illustration of our method, we applied it to somatic mutation data from 30 cancer types [8]. We applied the method to each cancer type separately to assess similarity of estimated signatures across cancer types. For each cancer type we selected the number of signatures *K* by fitting the model with increasing *K* and examining the log-likelihood, bootstrap errors, and correlation of membership parameters.The selected values of *K* are given in S2 Table. Also, we simply removed somatic mutations located in an intergenic region to include transcription strand biases as mutation features. Finally, we merged similar mutation signatures across different cancer types (when their Frobenius Distance were *<* 0.6).

Figs. 4 and 5 summarise the results. In total, we identified 27 mutation signatures. Many of these signatures show reassuring similarities with signatures identified in previous studies. However signatures from our independence model, because of its ability to effectively and parsimoniously deal with both *±*2 flanking base context and strand bias, are often more refined, highlighting additional details or features not previously evident. By comparing the composition of nucleotides and cancer types exhibiting the signatures with results of previous studies, we were able to associate many of the detected signatures with known mutational processes. In addition, as we reviewed these signatures and compared them with previous work, we noticed connections that, while not directly related to our new model, appear novel and noteworthy. The remainder of this section provides a comprehensive discussion of these findings.

**Figure 4.**
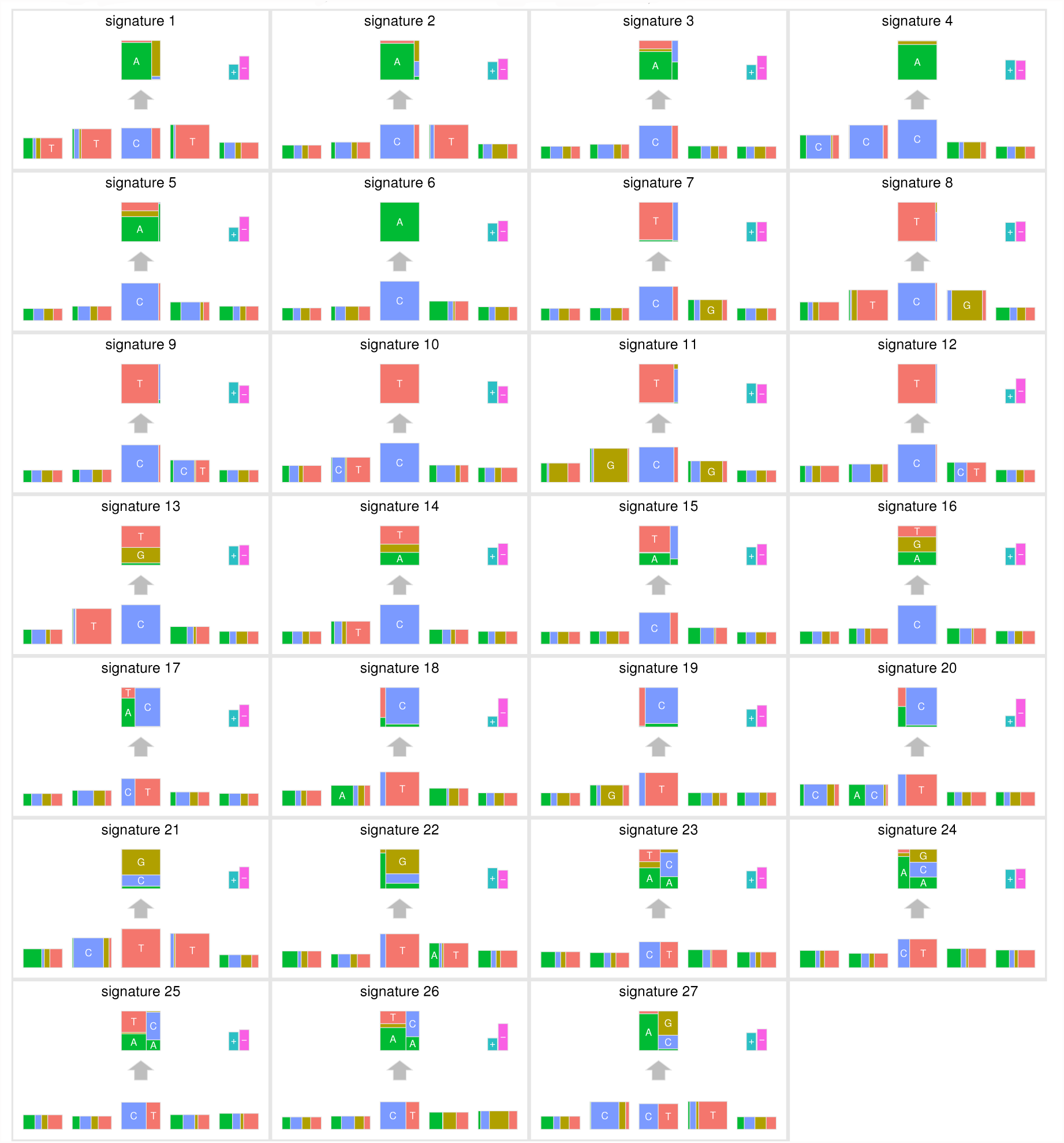
The summary of mutation signatures across 30 cancer types [8] obtained using the proposed method. Here, the substitution patterns and two 5’ and 3’ bases from the mutated sites are taken into account as mutation features. First, mutation signatures were estimated separately in each cancer type, and then similar signatures were merged (see text).

**Figure 5.**
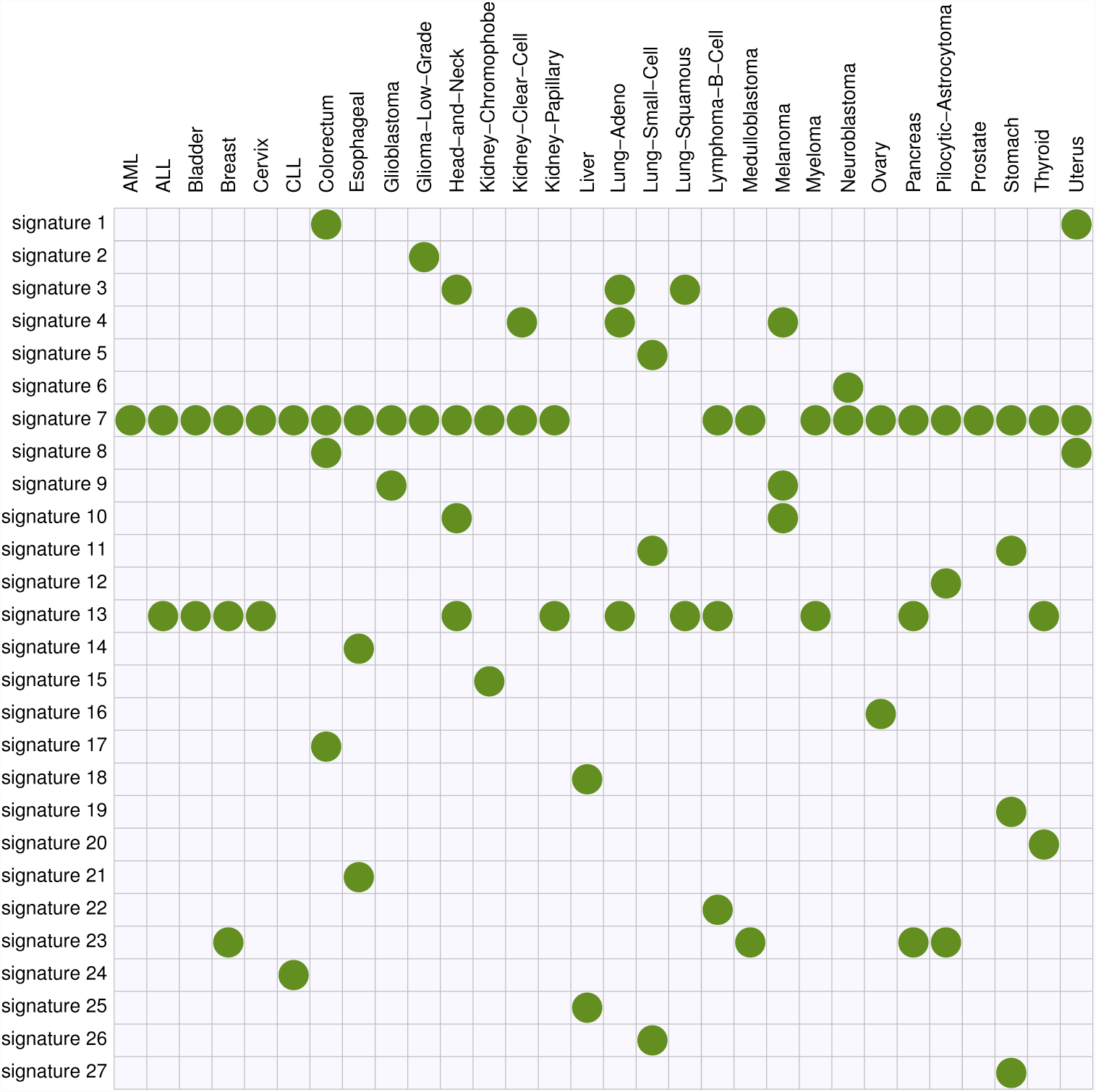
The summary of membership of each mutation signature across 30 cancer types obtained using the proposed method.

Signatures 1 and 8 (C *>* A at TpCpT and C *>* T at TpCpG, respectively) observed in colorectal and uterine cancers appear likely to be associated with deregulated activity of the error-prone polymerase Pol *ϵ* In previous analyses of these data [8], the signature for Pol *ϵ* dysfunction was represented by a single signature (their “signature 10”). In contrast our new approach uses two signatures. Since these signatures are highly correlated, and appear connected by a single biological mechansim, we certainly do not argue that inferring them as a single signature is “wrong”. However, splitting them into two signatures does help highlight certain features. Specifically, signature 1 shows a transcription strand bias whereas signature 8 does not, and this is true for both colorectal and uterus cancers (Figs. S2C S2D). This strand bias may be connected with the enrichment of C *>*A at TpCpT mutations in leading strands of replication forks observed by [19]. Although replication strand bias is different from transcription strand bias, these two biases may be connected through the fact that replication origins prefer transcription start sites [20].

These signatures also illustrate the ability of our model to help highlight sequence context effects beyond the immediate flanking bases. Specifically, both signatures 1 and 8 show an elevated frequency of the T base at position –2, and signature 1 also shows slightly elevated frequency of the T base at position +2 (Figs. 6B, 6C, S6C and S6D). A previous study of Pol *ϵ* [19] *found that a nonsense mutation R23X of TP53 is enriched in cancers with Pol ϵ* defects. In fact, the pattern of this mutation is C *>* T at TpTpCpGpA, closely matching signature 8. This illustrates that the inclusion of *±* 2 bases into signatures may be helpful for identifying underlying mechanisms.

**Figure 6.**
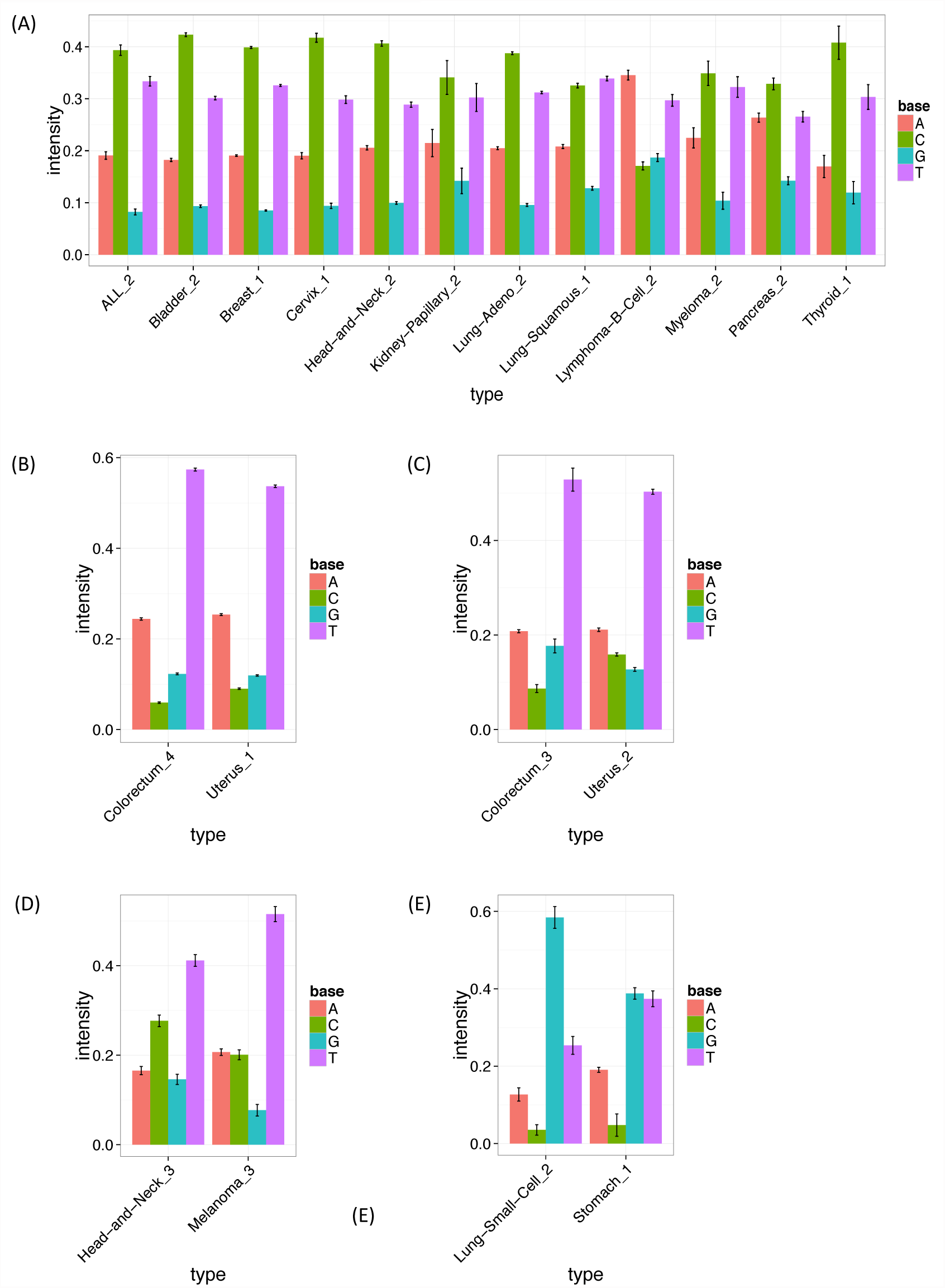
The estimated frequencies of bases at two 5’ to the mutated site for each cancer type. The bar heights show the estimated frequency for bases A, C, G and T at two 5’ to the mutated site. The error bars show bootstrapped standard errors. (A, B, C, D, E) The intensities of signature 13 (APOBEC signature), signature 1 (the first POL *ϵ* signature), signature 8 (the second POL ϵ signature), signature 10 (ultraviolet signature) and signature 11, respectively, at two 5’ to the mutated site. intensities at two 5’ to the mutated site.

Signature 2 (C *>* A at [CT]pCpT) is observed solely in low grade gliomas, and appears related to, but slightly different from, the signature previously detected in the same cancer types (“signature 14”, [9]). Indeed, the corresponding signature in the previous study shows very complex patterns (C *>* A at NpCpT or C *>* T at GpTpN). Further investigation revealed that this signature is driven by a single sample with an extremely high mutation rate (see Figs. S3A and S3B), and signature 2 disappeared when we removed this sample (see Fig. S4). It may be that the complex low-grade-glioma specific signature detected in the previous study is driven by the same single sample. We suggest that these signatures should be treated with caution until validated in additional samples.

Signature 4 (C *>* A at CpCpG) observed in kidney clear cell carcinomas, lung adenocarcinomas and melanomas seems to correspond to the “signature R2” detected in the same cancer types (plus lung squamous carcinomas) in [9] (see their Supplementary Figures). Again our analysis highlights additional contextual information, with a strikingly elevated frequency of base C at the –2 position (Figs. S5A, S5B and S5C). However, for each cancer type, only a few samples support this signature (see Figs. S5D, S5E and S5F), and the corresponding signature could not be validated in the previous study: most somatic mutations corresponding to that signature could not be validated by re-sequencing or visual inspection of BAM files using genomic viewers. Again, further investigation yielded a potential explanation for this finding: this signature largely matches that of a putative artifact caused by oxidation of DNA during acoustic shearing [21], and we conclude that this signature, and the corresponding signature in previous work, are likely artefactual. Although not of direct biological interest, identifying artefactual signatures could be helpful in removing false positive mutations.

Signature 13 (T *>* [AGT] at TpCpN sites) was observed in 12 cancer types, and is surely related to the activity of the APOBEC family. The 12 inferred signatures are highly consistent among cancer types except for B-cell lymphoma (see Fig. S2A), highlighting the robustness of our approach. Almost all of them show enrichment of A and T and depletion of G base at the –2 position (Fig. 6A and S2A), consistent with the UCUT data above and previous analyses [9]. The estimated transcribed strand specificities varied among cancer types, suggesting that there is not consistent strand-specificity in APOBEC signatures (and the observed variation may be due to estimation errors). Signatures 15 and 16 may also be related to APOBEC, although the estimated signatures are sufficiently different from 13 that they were not merged into a single cluster by our specified clustering criteria.

Signatures 10, 11, 12, 19 and 21 provide further examples of our method refining previously-observed signatures, highlighting strand biases and/or sequence context effects, particularly 2 bases upstream of the substitution. Signature 10 (C *>*T at [CT]pCpC) was observed in head and neck cancers and melanomas, and probably relates to ultraviolet light. Consistent strand specificities among the two cancer types (Fig. S2E) matches previous results [9], but our analysis additionally highlights elevated abundance of T at the –2 position (Figs. 6D and S2E). Signature 11 (C *>* T at GpCp[CG]) appears in small-cell lung cancers and stomach cancers, and seems to be the same as “signature 15” in the previous study, whose function remains unclear. Again our analysis highlights elevated abundance of G at the –2 position (Figs. 6E and S2F). Signatures 12 (C *>* T at [CG]pCp[CT]), 19 (T *>* C at GpTpN) and 21 (T *>* [CG] at CpTpT) observed in pilocytic astrocytomas, stomach cancers and oesophagus cancers, respectively, agree well with those detected in the same cancer types in the previous study [9]. However our analysis again refines these signatures, highlighting a strand bias in all three, and sequence context effects at the -2 position in Signatures 12 and 21.

One signature, Signature 20, appears not to match any signatures in the previous analysis [9] and represents a potentially novel signature. This signature (T *>* C at [AC]pTpN) is observed in thyroid cancers, and shows a very strong strand specificity, which could be due to transcription-coupled nucleotide excision repairs. This signature may have been too weak for previous methods to detect, perhaps because the mutation ratio of thyroid cancer is low, possibly reflecting improved sensitivity of our more parsimonious model.

The remaining signatures largely recapitulate previous results. Signature 3 and 5 (C *>* A at NpCpN) observed in head-and-neck cancers and three types of lung cancers are probably associated with tobacco smoking. The estimated signature in each cancer type shows higher mutation prevalence on the template strand (Fig. S2B), which is consistent with the previous study [2, 9]. Signature 6 (C *>* A at NpCp[AT]) observed in neuroblastomas matches the pattern detected in the same cancer type in the previous study. Signature 7 (C *>* T at NpCpG sites) was observed in 25 out of 30 cancer type, and arguably relates to deamination of 5-methyl-cytosine. Signature 9 (C *>* T at NpCp[CT]) was observed in melanomas and glioblastomas, and is probably associated with a chemotherapy drug, temozolomide. Signature 18 (T *>* C at ApTp[AG]) observed in liver cancers has been shown to be more common in Asian cases than in other ancestries [16], though the source of this signature is still not clear. In this signature, we observe a very strong strand specificity as shown in [9, 16], suggesting a possible role for transcription-coupled nucleotide excision repairs.

## Discussion

In this paper, we presented new methods for inferring and visualizing mutation signatures from multiple cancer samples. The new methods exploit simpler, more parsimonious, models for mutation signatures than existing methods. This improves stability of statistical estimation, and easily allows a wider range of contextual factors (e.g. more flanking bases) to be incorporated into the analysis. In addition, we provide a new intuitive way to visualize the inferred signatures.

We have also emphasised the connection between mutation signature detection, and the use of mixed-membership models in other fields, particularly admixture analysis and document clustering. This connection naturally raises the possibility of improving the proposed approach by learning from experiences in those other fields. For example, in admixture analysis, [22] found that the use of a correlated prior on allele frequencies improved sensitivity to detect population clusters; this suggests that it might be fruitful to consider a correlated prior distribution on signatures, to allow that some signatures - perhaps in different cancers - may be similar to one another (though not identical). More generally, introducing certain prior distributions or penalty terms, such as sparsity-promoting penalties [23,24] and determinantal point process priors [25, 26] could improve both accuracy and interpretation. Further, as the scale of cancer genome data becomes large, more sophisticated computational approaches for estimating parameters may become necessary. We can potentially borrow a number of computational techniques such as those using EM-algorithm [27, 28], sequential quadratic programming [29], Gibbs sampling [12, 30] and variational methods [13, 31, 32]. Finally, to address the problem of determining the number of signatures, it may be fruitful to extend the framework to the Hierarchical Dirichlet processes [33].

Although we have focused on point substitution mutations in this paper, many other types of mutations occur in cancer genomes, including insertions, deletions, double nucleotides substitutions, structural variations and copy number alterations [34, 35]. Our framework could incorporate these additional mutation types, by summarizing them using appropriate mutation features. In some cases, choice of appropriate features may need investigation. For example, longer deletions could be represented by the length of deletion and the adjacent bases; for short deletions (a few bases) it may be fruitful to include the actual deleted bases as part of the features.

We have detected a number of mutation signatures having transcription strand biases, which are naturally considered to be associated with transcription activities. Therefore, to further understand the effect of transcription activities on mutational mechanisms, we can include gene expression or RNA polymerase II occupancies to mutation features, so that the relationships of strand biases and transcription activities will be clarified.

Although we believe our new methods already provide useful gains compared with existing approaches, the methods are perhaps even more important for their future potential to incorporate other contextual data, including epigenetic data, into mutation signature analysis. This is important, because local mutation densities are closely related to a number of genomic and epigenetic factors, such as GC content, repeat sequences, chromatin accessibility and modifications, and replication timing [36–39]. A recent study found that epigenetic information in the cell types of origin of the corresponding tumors is the most predictive [40] for local mutation densities. A growing range of epigenetic data from many cell types are now available, and it will be interesting to integrate these epigenetic factors into mutation signature analysis to help understand how these epigenetic factors influence DNA damage and repair mechanisms. Our work here provides a straightforward way to do this: epigenetic data can be simply added as features to the mutation signature. We believe that the value and impact of our work, and specifically our proposed approach to modelling mutation signatures via independent features, will grow as more and more features are incorporated into the analysis.

## Methods

### Parameter Estimation

The parameters {***f***_*k,l*_} and {***q***_*i*_} must be estimated from the available mutation data {***x***_*i,j*_}. Here we adopt a simple approach that uses an EM-algorithm to maximise the likelihood.

Let *g*_*i,****m***_ denote the number of mutations in the *i*-th sample that have mutation feature vector ***m***. In the E step of the EM algorithm, we calculate values of auxiliary variables *θ*_*i,k,****m***_ defined as

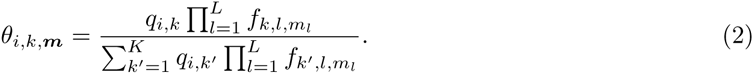

Then, in the M-step, we update the parameters {***f***_*k,l*_} and {*q*_*i,k*_} as

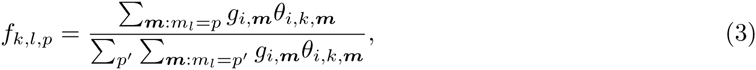

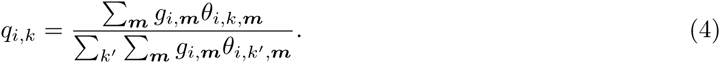

We use the R package SQUAREM [41] to accelerate convergence of this EM algorithm (SQUAREM implements a general approach to accelerate the convergence of any fixed-point iterative scheme such as an EM algorithm). To address potential problems with convergence to local minima, we apply the EM algorithm several times (10 times in this paper) using different initial points, and use the estimate with the largest log-likelihood. See S3 Text for the derivation of the above updating procedures.

### Background signatures

Here, we describe how the background mutation signature is obtained in the case where mutation features are the substitution patterns, the *±* 2 flanking bases, and the transcription strand. Since the majority of the data used in this paper is exome sequencing data, and since we consider transcription strand as a mutation feature, we use the exonic regions of the human genome reference sequence to obtain the background mutation signature. First we calculate the frequencies of 5-mers with transcription strands of the corresponding exon, where we take complement sequences and flip the strand for those whose central bases are A or G. Then, assuming alternated bases are equally likely from each central base C and T, the frequency of each mutation feature is derived directly from those of the 5-mers and transcription strands. Finally, the probability of each mutation feature is derived by normalizing each frequency to sum to one.

### Estimating standard errors

We use the non-parametric bootstrap [42] to calculate standard errors for parameter estimates. This involves resampling somatic mutations from the empirical distribution of the original data {***x***_*i,j*_} for each cancer genome. For each of 100 such bootstrap samples, we re-fitted the model, using parameters obtained for the original data as initial points. We then used sample standard errors of the inferred mutational signatures as estimates of parameter standard errors.

### Selecting the number of signatures

Determining an appropriate number of mutation signatures *K* is a challenging task. One approach is to utilize some statistical information criteria such as AIC [43] or BIC [44]. In the population structure problems, for example, the Bayesian deviance [12] and cross-validation [45] have been suggested. One previous study on mutation signature problems [10] utilized BIC. On the other hand, the problem of using these statistical information criteria is that most of them are based on the likelihood, where slight deviations between the specified probabilistic models and the reality sometimes leads to additional (possibly spurious) mutation signatures being selected to compensate for those deviations.

In this paper, instead of utilizing a statistical information criteria, we adopt the following strategy:

- After calculating the likelihood and standard errors of parameters for a range of *K*, the value of *K* is determined at the point where the likelihood is sufficiently high, and the standard errors are sufficiently low [9].
- When, for *k*_1_-th and *k*_2_-th mutation signatures, we could detect strong correlations between the estimated membership parameters for each cancer genome ((*q*_1,*k*_*1*__, *q*_2,*k*_*1*__, *…, q*_*I,k*_*1*__) and (*q*_1,*k*_*2*__, *q*_2,*k*_*2*__, *…, q*_*I,k*_*2*__)), and the two mutation signatures ({***F***_*k*_*1*__} and {***F***_*k*_*2*__}) show similar patterns, then this suggests that the method may have split one mutation signature into two. We choose *K* to be small enough that such pairs of mutation signatures do not occur.

These strategies are not claimed as optimal, but appeared to provide satisfactory results in our applications here. The development of automated and practical approaches for choosing *K* is a possible area for future development.

### Existing methods as a special case

Previous approaches to mutation signature modelling in [8, 9] are a special case of our framework. Specifically, they correspond to combining all possible combinations of mutation features into a single “meta-feature”, which takes *M*_1_ *× M*_2_ *×…× M*_*L*_ possible values. Thus, instead of having *L* features with ***M*** = (*M*_1_, *…, M*_*L*_), existing approaches have one feature with ***M*** = (*M*_1_ *×…× M*_*L*_) (see S1 Table). The resulting model allows for arbitrary distributions on the *M*_1_ ×…× *M*_*L*_ feature space, and we call the resulting model the “full model”. The full model can represent complicated dependencies in a single signature. For example, a situation where C *>* A is frequent at ApCpG sites and C *>* T is frequent at TpCpA sites could be represented with one signature. This may be desirable in some settings and not in others. However, when many mutation contextual factors are taken into account and the number of free parameters becomes huge, estimated results can be unstable and unreliable. Furthermore, there is a risk of over-interpreting the complex features of estimated signatures.

### Relationship with mixed-membership models

Our model is closely related to mixed-membership models that have been adopted in other applications, such as document classification and population structure inference problems. In this subsection, we outline these relationships, slightly abusing notation to contrast the relationships.

In the topic model [13, 27], which are a form of mixed-membership models frequently used in document classification problems, each document is assumed to have *K* different “topics” in varying proportions (***q***_*i*_ ∈ Δ^*K*^), where each topic is characterized by a word frequency (a multinomial distribution on a set of words *W* (***f***_*k*_ ∈ Δ^*W*^). And each word is assumed to be generated by one of *K* multinomial distributions (topics). The detailed generative process of the *j*-th word in the *i*-th document *x*_*i,j*_ is:

1. Generate the underlying topic for the *j*-th word: *z*_*i,j*_ *∼* Multinomial(***q***_*i*_), where *z*_*i,j*_ ∈ {1, *…, K}*.
2. Generate *x*_*i,j*_ *∼* Multinomial(***f*_*z*_*i,j*__**), where *x*_*i,j*_ ∈ {1, *…, W}*.

Actually our “full model” (*L* = 1) is essentially the same as a topic model.

On the other hand, in population structure inference problems [12, 46], each individual is assumed to be an admixture of *K* ancestries in varying proportions, where each ancestry is characterized by the allele frequency at each SNP locus. Each SNP genotype of an individual is assumed to be generated by the two step model: first, an ancestry (“population”) is chosen according to the admixture proportion for each individual, and then the SNP genotype is generated according to the allele frequency of the selected ancestry at that locus. The relationships among the mutation signature models, topic models and population structure models are summarized in Table 1.

**Table 1.**
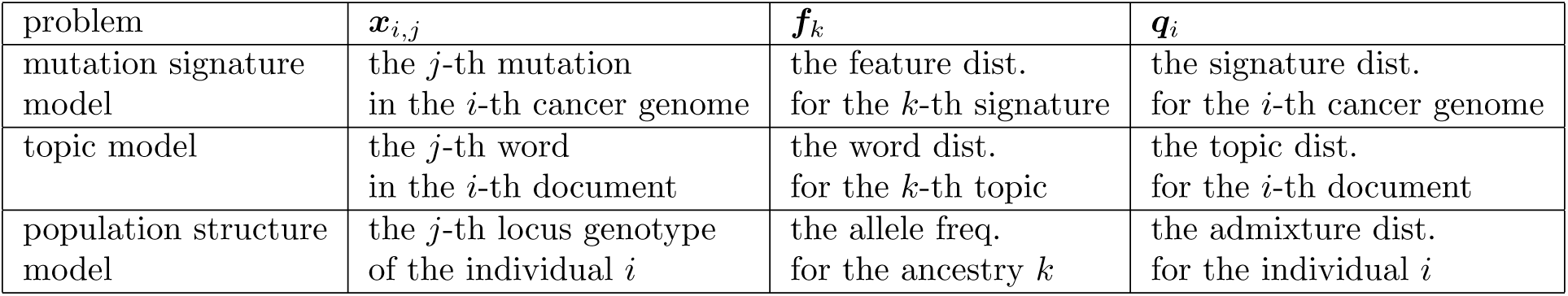
Relationships among mutation signature model, topic models, and population structure models.

As pointed out by [47], there is a close relationships between mixed-membership models and nonnegative matrix factorization, which has been successfully used in the previous studies for mutational signature problems [7–9]. In fact, the proposed method can be seen as non-negative matrix factorization with additional restrictions. See S4 Text for details of the relationship between the proposed approach and nonnegative matrix factorization.

## Supporting Information

**S1 Text**

**Experiments on synthetic data.**

**S2 Text**

**Experiment on UCUT data with the various number of mutation signatures.**

**S3 Text**

**Derivation of EM algorithm.**

**S4 Text**

**Relationship with nonnegative matrix factorization.**

**S1 Table**

**Example of representation for mutation patterns (substitution patterns and one 5’ and 3’ bases).** In the independent representation, the elements of vector show substitution patterns, 5’ adjacent bases and 3’ adjacent bases, respectively. For substitution pattens, 1 to 6 values are assigned to C*>*A, C*>*G, C*>*T, T*>*A, T*>*C and T*>*G in this order. For 5’ and 3’ adjacent bases, 1 to 4 values are assigned to A, C, G and T. Note that the original base is fixed to C or T to remove the redundancy of complement sequences.

**S2 Table**

**The number of mutation signatures selected for each cancer type for each cancer type in the Alexandrov et al. (2013) data.**

**S1 Figure**

**The frequencies of bases at two 5’ to the mutated site for the APOBEC mutation signatures obtained in UCUT data using the independent and full models.**

**S2 Figure**

**The list of several signatures extracted in each cancer type in the Alexandrov et al. (2013) data.** (A) APOBEC signatures (signature 13) obtained in each cancer type. (B) Smoking signature in each cancer type. (C) The first Pol *ϵ* signature (signature 1) in each cancer type. (D) The second Pol *ϵ* signature (signature 8) in each cancer type. (E) The ultraviolet signature i(signature 10) n each cancer type. (F) Unknown signature (signature 11) obtained in lung small cell carcinomas and stomach cancers.

**S3 Figure**

**The estimated membership parameters of low grade gliomas.** (A, B) Estimated membership parameter by the proposed method in normal and log scale. We have selected top 100 cancer samples according to the number of mutation. The height of bar shows (the logarithm of) the number of mutations for each sample, and the ratio of colored division shows the ratio of estimated membership parameters for each signature and sample. The low grade glioma specific signature detected by the proposed method is the signature 2. We can see that the mutations corresponding to signature 2 is mostly from the sample with an extremely high mutation rate.

**S4 Figure**

**The signatures obtained for the original data and for the data without the hyper-mutated case.** (A) The result for the original data (*K* = 3). The first (from the left) signature seems to be one from deamination of 5-methyl-cytosine. The second signature is the low grade glioma signature. (B, C, D, E) The result for the data without the hyper-mutated sample for *K* = 2, 3, 4 and 5, respectively. Although the signature related to deamination of 5-methyl-cytosine remained, low grade glioma specific signature could not be observed.

**S5 Figure**

**The putative oxidative artifact signatures and membership parameters estimated for each cancer.** (A) The second signature detected in kidney clear cell carcinomas. (B) The first signature detected in lung adenocarcinomas. (C) The first signature detected in melanomas (D, E, F) Estimated membership parameter for kidney clear cell carcinomas, lung adenocarcinomas and melanomas, respectively. For each cancer type, We have selected top 100 cancer samples according to the number of mutation. The height of bar shows the number of mutations for each sample, and the ratio of colored division shows the ratio of estimated membership parameters for each signature and sample. We can see that the signature corresponding to putative oxidative artifacts concentrates on a small number of samples.

## Acknowledgments

We thank to Dr. Daichi Mochihashi for helpful discussion and comments on an earlier version of the method; Dr. Hiromichi Suzuki for many helpful feedback on the R packages; Dr. Ziyue Gao for reading the manuscript and providing many helpful comments.

This project was deeply influenced by the broader intellectual and academic environment surrounding Matthew Stephens’s Laboratory, when the first author stayed at the University of Chicago as a visiting scholar. The first author thanks Dr. John Novembre, Dr. Jacob Degner and all the members of Matthew Stephens and John Novembre Laboratories for helpful discussion and comments.

## References

1. Stratton MR, Campbell PJ, Futreal PA. The cancer genome. Nature. 2009;458(7239):719–724.

2. Pfeifer GP, Denissenko MF, Olivier M, Tretyakova N, Hecht SS, Hainaut P. Tobacco smoke carcinogens, DNA damage and p53 mutations in smoking-associated cancers. Oncogene. 2002 Oct;21(48):7435–7451.

3. Pfeifer GP, You YH, Besaratinia A. Mutations induced by ultraviolet light. Mutat Res. 2005 Apr;571(1-2):19–31.

4. Burns MB, Lackey L, Carpenter MA, Rathore A, Land AM, Leonard B, et al. APOBEC3B is an enzymatic source of mutation in breast cancer. Nature. 2013 Feb;494(7437):366–370.

5. Burns MB, Temiz NA, Harris RS. Evidence for APOBEC3B mutagenesis in multiple human cancers. Nat Genet. 2013 Sep;45(9):977–983.

6. Roberts SA, Lawrence MS, Klimczak LJ, Grimm SA, Fargo D, Stojanov P, et al. An APOBEC cytidine deaminase mutagenesis pattern is widespread in human cancers. Nat Genet. 2013 Sep;45(9):970–976.

7. Nik-Zainal S, Alexandrov LB, Wedge DC, Van Loo P, Greenman CD, Raine K, et al. Mutational processes molding the genomes of 21 breast cancers. Cell. 2012 May;149(5):979–993.

8. Alexandrov LB, Nik-Zainal S, Wedge DC, Aparicio SA, Behjati S, Biankin AV, et al. Signatures of mutational processes in human cancer. Nature. 2013 Aug;500(7463):415–421.

9. Alexandrov LB, Nik-Zainal S, Wedge DC, Campbell PJ, Stratton MR. Deciphering signatures of mutational processes operative in human cancer. Cell Rep. 2013 Jan;3(1):246–259.

10. Fischer A, Illingworth CJ, Campbell PJ, Mustonen V. EMu: probabilistic inference of mutational processes and their localization in the cancer genome. Genome Biol. 2013 Apr;14(4):R39.

11. Krawczak M, Ball EV, Cooper DN. Neighboring-nucleotide effects on the rates of germ-line single-base-pair substitution in human genes. Am J Hum Genet. 1998 Aug;63(2):474–488.

12. Pritchard JK, Stephens M, Donnelly P. Inference of population structure using multilocus genotype data. Genetics. 2000 Jun;155(2):945–959.

13. Blei DM, Ng AY, Jordan MI. Latent Dirichlet Allocation. J Mach Learn Res. 2003 Mar;3:993–1022. Available from: http://dl.acm.org/citation.cfm?id=944919.944937.

14. Eddelbuettel D, François R, Allaire J, Chambers J, Bates D, Ushey K. Rcpp: Seamless R and C++ integration. Journal of Statistical Software. 2011;40(8):1–18.

15. Schneider TD, Stephens RM. Sequence logos: a new way to display consensus sequences. Nucleic Acids Res. 1990 Oct;18(20):6097–6100.

16. Totoki Y, Tatsuno K, Covington KR, Ueda H, Creighton CJ, Kato M, et al. Trans-ancestry mutational landscape of hepatocellular carcinoma genomes. Nat Genet. 2014 Dec;46(12):1267–1273.

17. Rrnyi A. On measures of entropy and information. In: Fourth Berkeley symposium on mathematical statistics and probability. vol. 1; 1961. p. 547–561.

18. Hoang ML, Chen CH, Sidorenko VS, He J, Dickman KG, Yun BH, et al. Mutational signature of aristolochic acid exposure as revealed by whole-exome sequencing. Sci Transl Med. 2013 Aug;5(197):197ra102.

19. Shinbrot E, Henninger EE, Weinhold N, Covington KR, Goksenin AY, Schultz N, et al. Exonuclease mutations in DNA polymerase epsilon reveal replication strand specific mutation patterns and human origins of replication. Genome Res. 2014 Nov;24(11):1740–1750.

20. Dellino GI, Cittaro D, Piccioni R, Luzi L, Banfi S, Segalla S, et al. Genome-wide mapping of human DNA-replication origins: levels of transcription at ORC1 sites regulate origin selection and replication timing. Genome Res. 2013 Jan;23(1):1–11.

21. Costello M, Pugh TJ, Fennell TJ, Stewart C, Lichtenstein L, Meldrim JC, et al. Discovery and characterization of artifactual mutations in deep coverage targeted capture sequencing data due to oxidative DNA damage during sample preparation. Nucleic Acids Res. 2013 Apr;41(6):e67.

22. Falush D, Stephens M, Pritchard JK. Inference of population structure using multilocus genotype data: linked loci and correlated allele frequencies. Genetics. 2003 Aug;164(4):1567–1587.

23. Hoyer PO. Non-negative matrix factorization with sparseness constraints. The Journal of Machine Learning Research. 2004;5:1457–1469.

24. Engelhardt BE, Stephens M. Analysis of population structure: a unifying framework and novel methods based on sparse factor analysis. PLoS genetics. 2010;6(9):e1001117.

25. Kulesza A, Taskar B. Determinantal point processes for machine learning. arXiv preprint arXiv:12076083. 2012;.

26. Kwok JT, Adams RP. Priors for diversity in generative latent variable models. In: Advances in Neural Information Processing Systems; 2012. p. 2996–3004.

27. Hofmann T. Probabilistic Latent Semantic Indexing. In: Proceedings of the 22Nd Annual International ACM SIGIR Conference on Research and Development in Information Retrieval. SIGIR ’99. New York, NY, USA: ACM; 1999. p. 50–57. Available from: http://doi.acm.org/10.1145/312624.312649.

28. Tang H, Peng J, Wang P, Risch NJ. Estimation of individual admixture: analytical and study design considerations. Genetic epidemiology. 2005;28(4):289–301.

29. Zhou H, Alexander D, Lange K. A quasi-Newton acceleration for high-dimensional optimization algorithms. Statistics and computing. 2011;21(2):261–273.

30. Griffiths TL, Steyvers M. Finding scientific topics. Proc Natl Acad Sci USA. 2004 Apr;101 Suppl 1:5228–5235.

31. Teh YW, Newman D, Welling M. A collapsed variational Bayesian inference algorithm for latent Dirichlet allocation. In: Advances in neural information processing systems; 2006. p. 1353–1360.

32. Raj A, Stephens M, Pritchard JK. Variational Inference of Population Structure in Large SNP Datasets. Genetics. 2014;p. genetics–114.

33. Teh YW, Jordan MI, Beal MJ, Blei DM. Hierarchical dirichlet processes. Journal of the american statistical association. 2006;101(476).

34. Meyerson M, Gabriel S, Getz G. Advances in understanding cancer genomes through second-generation sequencing. Nature Reviews Genetics. 2010;11(10):685–696.

35. Helleday T, Eshtad S, Nik-Zainal S. Mechanisms underlying mutational signatures in human cancers. Nature Reviews Genetics. 2014;15(9):585–598.

36. Schuster-Bockler B, Lehner B. Chromatin organization is a major influence on regional mutation rates in human cancer cells. Nature. 2012 Aug;488(7412):504–507.

37. Hodgkinson A, Chen Y, Eyre-Walker A. The large-scale distribution of somatic mutations in cancer genomes. Hum Mutat. 2012 Jan;33(1):136–143.

38. Liu L, De S, Michor F. DNA replication timing and higher-order nuclear organization determine single-nucleotide substitution patterns in cancer genomes. Nat Commun. 2013;4:1502.

39. Lawrence MS, Stojanov P, Polak P, Kryukov GV, Cibulskis K, Sivachenko A, et al. Mutational heterogeneity in cancer and the search for new cancer-associated genes. Nature. 2013 Jul;499(7457):214–218.

40. Polak P, Karli R, Koren A, Thurman R, Sandstrom R, Lawrence MS, et al. Cell-of-origin chromatin organization shapes the mutational landscape of cancer. Nature. 2015 Feb;518(7539):360–364.

41. Varadhan R, Roland C. Simple and globally convergent methods for accelerating the convergence of any EM algorithm. Scandinavian Journal of Statistics. 2008;35(2):335–353.

42. Efron B, Tibshirani RJ. An introduction to the bootstrap. CRC Press; 1994.

43. Akaike H. A new look at the statistical model identification. Automatic Control, IEEE Transactions on. 1974;19(6):716–723.

44. Schwarz G, et al. Estimating the dimension of a model. The annals of statistics. 1978;6(2):461–464.

45. Alexander DH, Lange K. Enhancements to the ADMIXTURE algorithm for individual ancestry estimation. BMC bioinformatics. 2011;12(1):246.

46. Alexander DH, Novembre J, Lange K. Fast model-based estimation of ancestry in unrelated individuals. Genome Res. 2009 Sep;19(9):1655–1664.

47. Ding C, Li T, Peng W. On the equivalence between non-negative matrix factorization and probabilistic latent semantic indexing. Computational Statistics & Data Analysis. 2008;52(8):3913– 3927.

